# To block or not to block: the adaptive manipulation of plague transmission

**DOI:** 10.1101/392514

**Authors:** S. Gandon, L. Heitzmann, F. Sebbane

## Abstract

The ability of the agent of plague, *Yersinia pestis*, to form a biofilm blocking the gut of the flea has been considered to be a key evolutionary step in maintaining flea-borne transmission. However, blockage decreases dramatically the life expectancy of fleas, challenging the adaptive nature of blockage. Here we develop an epidemiological model of plague that accounts for its different transmission routes, as well as the within-host competition taking place between bacteria within the flea vector. We use this theoretical framework to identify the environmental conditions promoting the evolution of blockage. We also show that blockage is favored at the onset of an epidemic, and that the frequencies of bacterial strains exhibiting different strategies of blockage can fluctuate in seasonal environments. This analysis quantifies the contribution of different transmission routes in plague and makes testable predictions on the adaptive nature of blockage.

**Significance statement:** Plague transmission relies on the ability of infected fleas to inoculate *Y. pestis* bacteria to vertebrate hosts. The production of a biofilm by the bacteria blocks the forgut of the flea and increases infectivity. But the adaptive nature of blockage remains controversial because it has a massive survival cost on the infected fleas and reduces dramatically the length of the infection: an extreme form of the classical virulence-transmission tradeoff. Here we develop a comprehensive model of the multiple routes of plague transmission, we determine when blockage can be viewed as an adaptive manipulation of its flea vector and we generate several testable predictions on the evolution of plague in both endemic and epidemic situations.

---

*Yersinia pestis* is the bacterium that caused 3 plague pandemics and had a profound effect on human history (1). A combination of comparative genomics analyses and experimental studies have unvealed the different evolutionary steps leading to the emergence and the spread of one of the deadliest human pathogen. *Y. pestis* recently emerged from *Yersinia pseudotuberculosis*, a food- and waterborne enteric pathogen causing a benign disease of the digestive tract in humans (2–5). Only a handful of genetic events, including acquisition of genes by horizontal transfer and loss of functional genes, led to the production of flea-borne transmission of plague (3,4,6,7). Notably, the horizontal acquisition of the *Yersinia* murin toxin gene (*ymt*) that protects from a bacteriolytic agent generated during the digestion of the blood meal has been essential to colonize the flea’s midgut and foregut (8). Truncation of the urease accessory protein UreD due to the insertion of a single nucleotide in the *ureD* locus (pseudogenization) reduced the toxicity of the ancestral strain, thereby prolonging the duration of infection in the vector (9,10). Lastly, a series of pseudogenization which led to the loss of the functional accessory regulatory protein RcsA and of two phospodisterases (PDE) unlocked the pre-existing capability of the ancestral strain to form a biofilm thanks to the *hmsHFRS* operon, enabling persistent colonization of the proventriculus and ultimately blockage of flea’s gut (7,11).

When the proventriculus of the flea is blocked the biofilm prevents the incoming blood to enter the midgut. The blood meal is contaminated upon contact with the bacterial mass, and is regurgitated at the flea-bite site, leading to transmission of plague (12). Another consequence of the blockage is an increase in the biting rate as the flea starves to death. Therefore, blockage is often viewed as a key adaptation of *Y. pestis* because it boosts bacterial transmission by increasing both infectivity (the number of bacteria inoculated in a new host) and the biting rate of infected fleas (7,11). Yet, the adaptive nature of blockage is challenged by the fact that it drastically increases the mortality rate of the flea (7,11). Besides, a combination of experimental observations and empirical studies suggest that other routes of transmission may be involved in plague epidemics (13–16,6,17,18,4). In particular, some flea transmission may also occur in an early-phase of the infection of unblocked flea. In other words, blockage may be viewed as a by-product of the colonization of the foregut but not as an adaptive manipulation of the biting rate of its insect vector.

To evaluate the relative importance of blockage on plague transmission we first develop a theoretical framework that accounts for the multiple transmission routes of *Y. pestis* (Figure 1). In a second step, we use this theoretical framework to study the evolution of the propensity to block the flea. To analyse pathogen evolution we study the competition between bacterial strains with varying blockage strategies. This competition takes place at a between-host level when bacteria are trying to infect new hosts. But bacteria may also compete within-host when, for instance, a flea is coinfected by different strains after feeding on two infected hosts. We derive threshold conditions allowing the invasion of a mutant strain with a specific blockage strategy in a stable environment. We also analyse the evolution of the plague during epidemics and show how bacteria with different rates of blockage can fluctuate in a seasonal environment.

**Figure 1:**
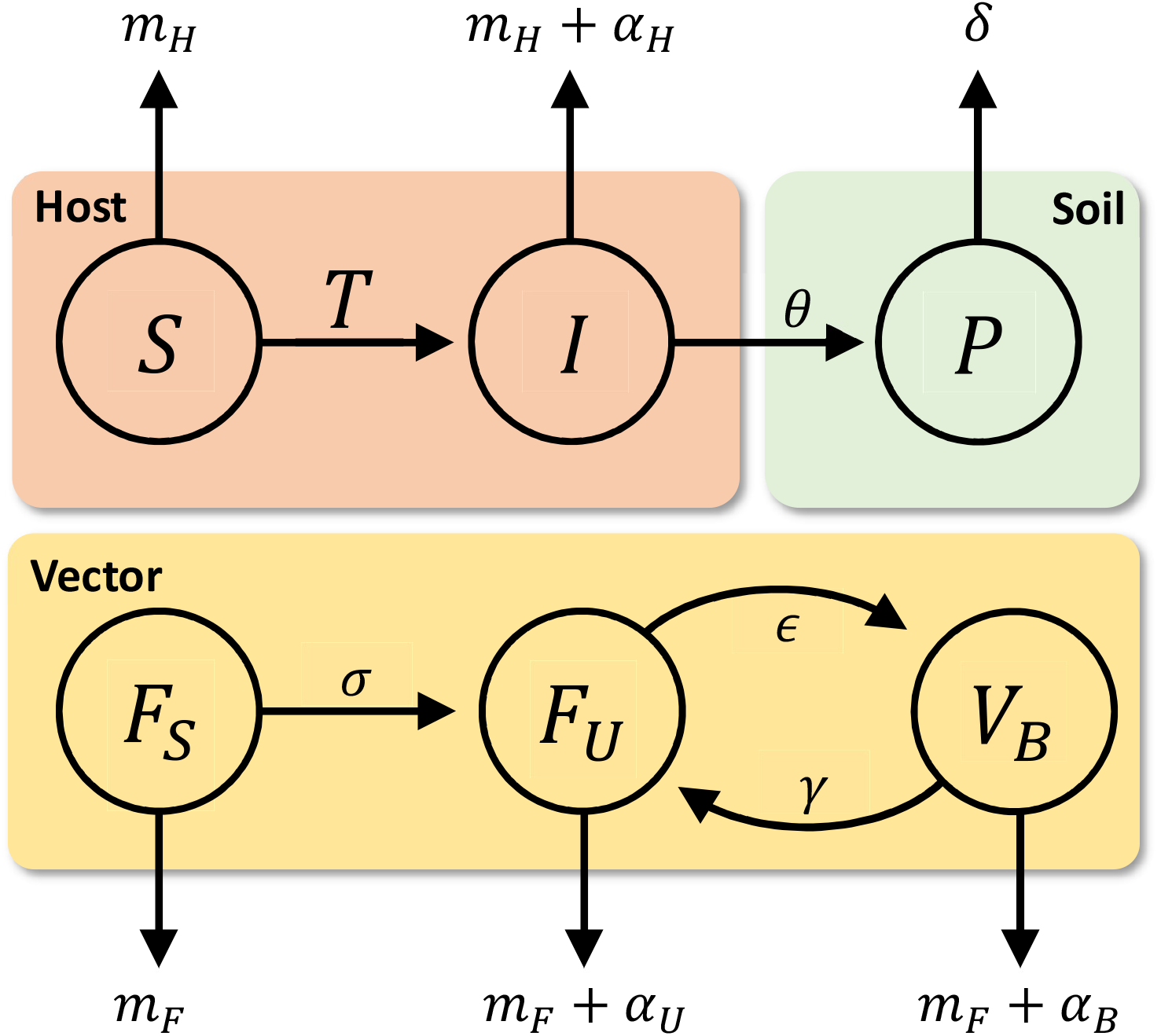
The multiple routes of plague transmission. Our model accounts for the circulation of *Y. pestis* in three different habitats: (1) a vertebrate host, (2) the vector and (3) the soil. The rate at which uninfected hosts become infected is determined by the sum of the force of infection from the different compartments of this system (see description of life cycle in the main text): *T* = *β_H_I* + *β_P_P* + *β_U_F_U_* + *β_B_F_B_*.

## The model

*Y. pestis* bacteria can live and/or persist in three different habitats: (i) a vertebrate host, (ii) a flea and (iii) the soil. Our epidemiological model accounts for the complex life-cycle of Y. *pestis* through these three different compartments of the environment (**Figure 1**). For the sake of simplicity we assume that the densities of both the vertebrate host (the host) and the flea (the vector) are constant and equal to *N_H_* and *N_F_*, respectively (we will relax this assumption later on in the analysis). The natural mortality rates of hosts and vectors are *m_H_* and *m_F_*, respectively. Because we are interested in plague evolution we assume that multiple bacterial strains can circulate. We note *P_i_* the density of the free-living stage of the strain *i*, and *I_i_* the density of hosts infected with the strain *i*. After feeding on a host infected with strain *i*, the infected flea is assumed to be “unblocked” state (state *F_U,i_*). Infectious fleas can become “blocked” (state *V_B,i_*) and the transition between the “unblocked” and the “blocked” states occurs at a rate *ϵ_i_* (the rate of blockage), which is assumed to vary among different strains of *Y. pestis*. We also assume that blocked fleas can become unblocked (return to the state *F_U,i_*) at a constant rate *γ*. Infection increases the mortality of the host (*α_H_*), and the mortality of both the blocked and the unblocked fleas (*α_B_* and *α_U_*, respectively). It is important to note that blockage has a major impact on flea survival (*α_B_* > *α_U_*) (11,7). Hence, bacterial strains that promote blockage are associated with higher virulence in the flea because blockage decreases survival. Note, however, that once the infected flea is blocked (or unblocked) all the strains have the same mortality rates. The host can acquire the infection horizontally from other infected hosts at a rate *β_H_I_i_*, from the propagules in a contaminated environment at a rate *β_P_P_i_* and from the infected vectors at rates *β_U_F_U,i_* and *β_B_F_B,i_*. The parameters *β_H_*, *β_P_*, *β_U_* and *β_B_* modulate the relative importance of these four different routes of transmission. Crucially, experimental studies have demonstrated that blockage increases the infectiousness of fleas and thus *β_B_* > *β_U_* (11,19,20,7). This life cycle can be summarized in the following system of equations (see **Table S1** for the definition of all the parameter of this model):

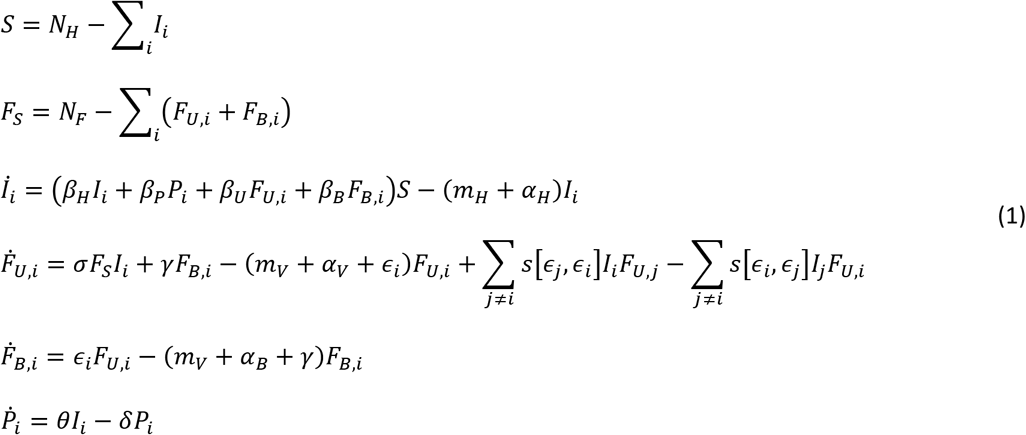

The above model accounts also for the competition taking place between bacterial strains in the early stage of the infection (i.e. in unblocked fleas). Indeed, when an unblocked flea infected with strain *i* feeds on a host infected by strain *j* the superinfection function *s*[*ϵ_i_, ϵ_j_*] determines the probability that strain *i* is replaced by strain *j*. We assume that the competitivity of the bacteria may be associated with the propensity to form biofilms and to block the flea. We used the following function to model superinfection:

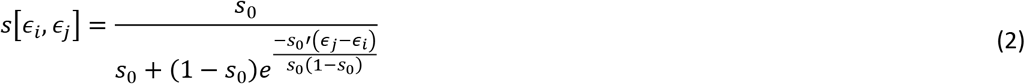

where *s*_0_ = *s*[*ϵ_i_, ϵ_i_*] is the value of the probability of superinfection at the origin (when both strains have the same value of *ϵ*) and 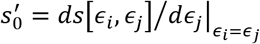 is the slope of the superinfection function at the origin (**Figure S1**).

Note that we neglect the possibility that competition may occur in blocked fleas and in vertebrate hosts because the bacterial density reached in blocked fleas and in infected hosts hampers invasion by new strains. This is arguably a very simplified view of the way within-host competition among bacterial strains may occur in this system. Yet, as we will see below, the simplicity of this model shows the potential implications of within-host competition on plague evolution and leads to novel adaptive hypothesis for the evolution of blockage.

## Epidemiology and evolution in a stable environment

First, we focus on a scenario where the population of the bacteria is monomorphic and all the parameters of the model are constant. The basic reproduction ratio *R*_0_ of the pathogen is given by (see **Appendix**):

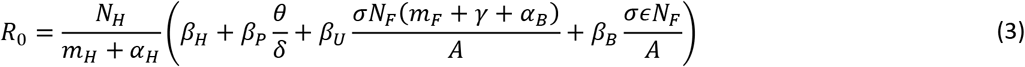

with *A* = *m_F_*(*m_F_* + *γ* + *ϵ*) + *α_U_*(*m_F_* + *γ*) + *α_B_*(*m_F_* + *α_U_* + *ϵ*). The above expression is useful to identify the relative importance of the different routes of transmission on the epidemiology of plague. Indeed, each term in the parenthesis are associated with the contribution of each of the 4 different routes of transmission to *R*_0_ : (i) direct horizontal transmission, (ii) transmission via propagules, (iii) transmission via unblocked fleas, (iv) transmission via blocked fleas.

This expression is also particularly useful to identify the conditions promoting the ability of the pathogen to trigger an epidemic in an uninfected host population where *S* = *N_H_* and *F_S_* = *N_F_*. When *R*_0_ > 1, the pathogen can invade the host population and the system reaches an endemic equilibrium where the host, the vector and the pathogen can coexist (the notation 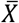 is used to refer to the equilibrium density of the variable *X* at this endemic equilibrium). Numerical exploration of the system (1) revealed that this endemic equilibrium was always locally stable.

In the following, we study the long-term evolutionary dynamics of plague using the classical formalism of Adaptive Dynamics, where mutation rate is assumed to be low which allows decoupling evolutionary and epidemiological dynamics (21–24). To study plague evolution we derive the invasion fitness per-generation of a *mutant* strain which has the strategy *ϵ_m_*, at the endemic equilibrium set by a resident population of the pathogen which has the strategy *ϵ* (25) (**Appendix**):

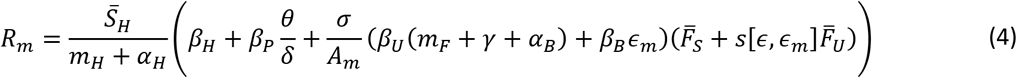

with: *A_m_* = *m_F_*(*m_F_* + *γ* + *ϵ_m_*) + *α_U_*(*m_F_* + *γ*) + *α_B_*(*m_F_* + *α_U_* + *ϵ_m_*) + *σs*[*ϵ_m_, ϵ*]*Ī*(*m_F_* + *γ* + *α_B_*). The mutant will invade the resident population if *R_m_* > 1 and this invasion fitness can be used to derive the gradient of selection on blockage at the endemic equilibrium (i.e. 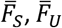 and 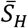) set by the resident strategy.

We used this invasion fitness to identify the conditions leading the evolution of higher rates of blockage (**Appendix**). In particular, under the assumption that the superinfection function is constant and equal to *s*_0_, we find that higher rates of blockage are selected for when:

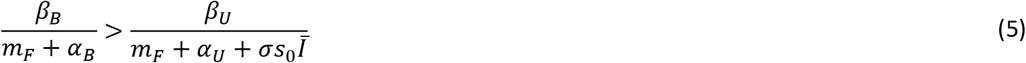

Hence, in spite of the complexity of the life cycle, the evolution of blockage boils down to a very simple condition that does not depend on the other routes of transmission. The left and the right hand sides of (5) measure of the relative quality of blocked and unblocked fleas, respectively. The quality of a vector depends on the instantaneous rate of transmission (*β_B_* and *β_U_*) but also the duration of the infection which is modulated by the mortality rates (*m_F_, α_U_* and *α_B_*) as well as the rate of superinfection in unblocked fleas (*σs*_0_*Ī*). Blockage evolves whenever the blocked fleas are better vectors than unblocked fleas. When condition (5) is satisfied blockage evolves to maximal values. In contrast, when condition (5) is not satisfied, blockage does not evolve and the evolutionary stable strategy is *ϵ*\* = 0.

The invasion condition can also be used to determine the conditions favoring the evolution of blockage when the probability of superinfection depends on the investment in blockage of the competing strains (i.e., *s*_0_′ ≠ 0). For instance, under the simplifying assumption that the resident strain does not block (*ϵ* = 0) the condition for the invasion of a mutant strain that blocks the flea is:

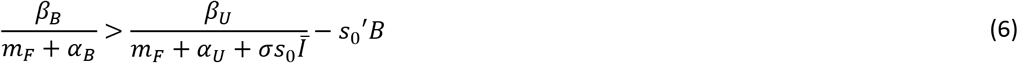

where 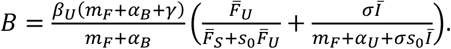

The above condition shows that if the ability to block the flea is associated with a higher competitive ability of the bacteria (i.e., 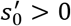), blockage can evolve more readily. In contrast, if the production of a biofilm is costly and induces a lower competitive ability (i.e., 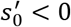), it is more difficult to evolve blockage. Besides, adding a cost on biofilm production allows some intermediate blockage strategy to be evolutionary stable (**Figure S2**).

## Evolution in a fluctuating environment

Because plague dynamics is often characterized by dramatic temporal fluctuations (26,27) we examined the evolution of blockage away from the endemic equilibrium. Numerical simulations show that, at the onset of an epidemic a mutant strain with a higher ability to block the flea can increase in frequency (**Figure 2**) even if this blockage strategy does not verify conditions (5) or (6). To understand pathogen evolution during this transient phase of the epidemics it is important to track both the *frequency* of the different strains and the *densities* of the pathogen in the different compartments of the model (28–31). In the following, we derive the dynamics of the frequencies 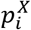, of the strain *i* in the compartment *X*:

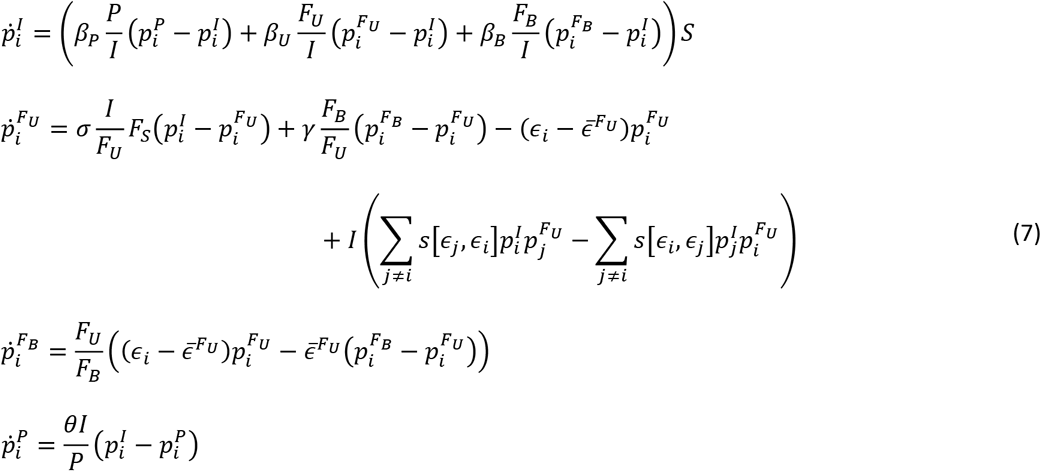

where 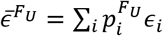 is the average value of blockage in unblocked fleas.

**Figure 2:**
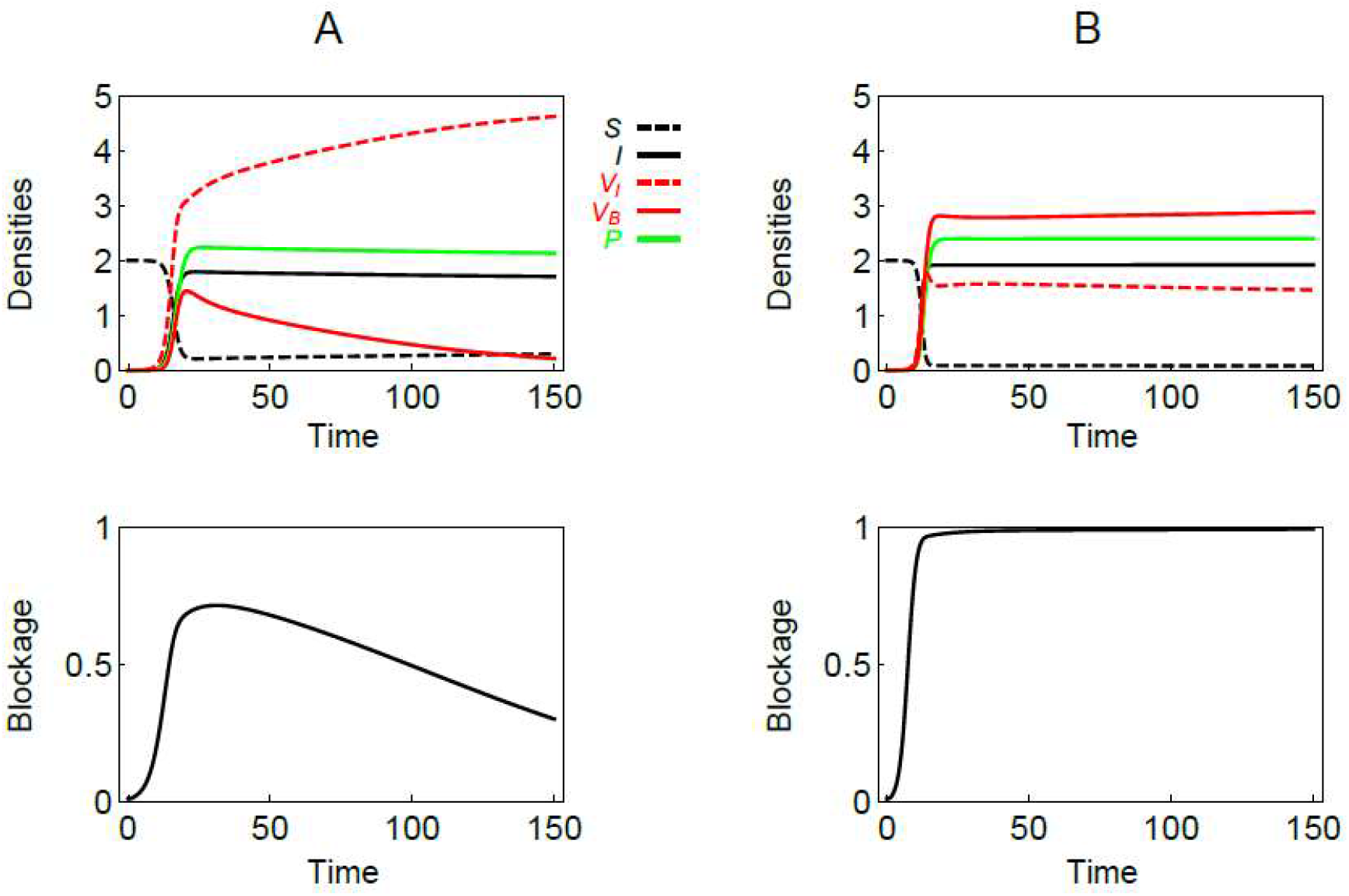
Epidemiology and evolution of plague during an epidemic. We present the epidemiological dynamics and the evolutionary dynamics in the absence of superinfection. The top figures show the dynamics of the densities of the different compartments of the model during an epidemic. The bottom figures show the dynamics of the mean value of the blockage strategy. We allow competition between two strains with very different blockage strategy (i.e. *ϵ* = 0 or 1). In panel (A) *β_B_* = 0.3 and blockage is maladaptive according to condition (5) (i.e.,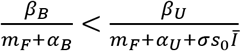 but blockage is selected for at the beginning of the epidemic. In panel (B) *β_B_* = 0.7 and blockage is adaptive according to condition (5) (i.e., 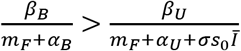. Other parameter values (see **Appendix** for more details about the simulation procedure): *N_H_* = 2, *N_F_* = 5, *γ* = 0.2, *θ* = 1, *σ* = 0.5, *δ* = 0.8, *m_H_* = 0.002, *m_F_* = 0.01, *α_H_* = 0.1, *α_U_* = 0.01, *α_B_* = 0.2, *β_H_* = 0.1, *β_P_* = 0.06, *β_U_* = 0.05.

Focusing on the dynamics of mutant frequency is particularly useful to understand the interplay between epidemiology and evolution. For instance, let us focus on the scenario where two bacterial strains compete: a mutant strain that blocks the fleas at a rate *ϵ_m_* and a resident strain that never blocks the fleas. In this case 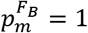 because only the mutant can block the fleas. If we neglect superinfections and assume the initial frequency of the mutant is low, the above dynamical system reduces to:

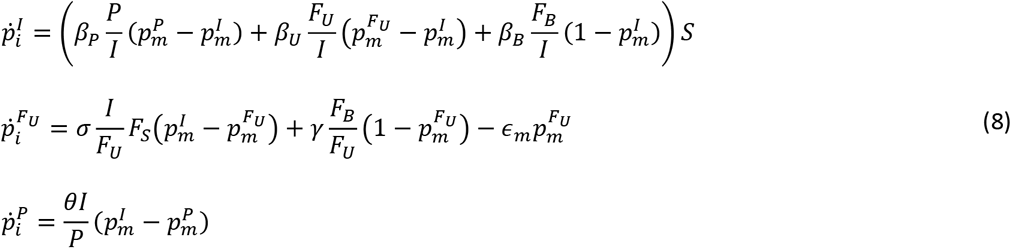

Initially, the mutant frequency is expected to be low in all the 3 other compartments of the model (*I, F_U_* and *P*) which yields the following approximation for the change in mutant frequency in the infected host compartment: 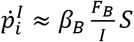. This indicates that the mutant frequency is initially increasing in the infected host compartment. This initial increase occurs even if the mutant is ultimately selected against (**Figure 2**). This transient selection for the mutant is due to the fitness benefit associated with higher transmission rates when there are a lot of susceptible hosts around (30,31).

If we impose periodic fluctuations in the densities of the host and the vector (e.g. induced by the seasonality of the environment), we observe periodic fluctuations of the incidence of the disease across time. These fluctuations maintain the pathogen away from the endemic equilibrium and can favor different blockage strategies in different phases of these recurrent epidemics. More blockage is selected for at the onset of the epidemics and it is selected against when the epidemics is fading away (**Figure 3**).

**Figure 3:**
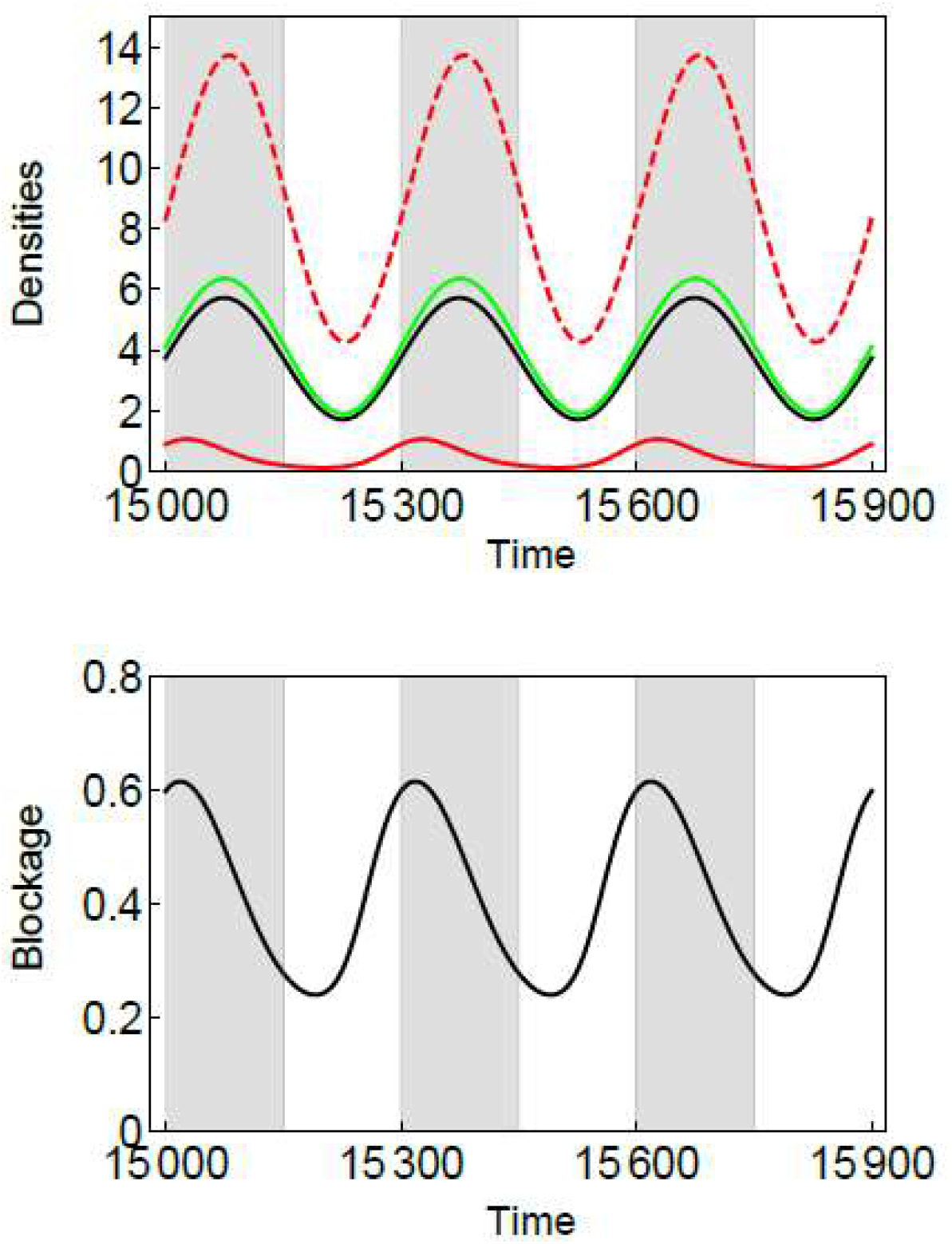
Epidemiology and evolution of plague in a seasonal environment. We allow the densities *N_H_* and *N_F_* to fluctuate periodically with the function *f*(*t*) = 1 + *Sin*(2*πt*/*T*), where *T* = 300 (the shaded area indicates time when *f*(*t*) > 1). We allow competition between two strains with very different blockage strategy (i.e. *ϵ* = 0 or 0.8). The two strains coexist but their relative frequencies fluctuate with the variations of the abundance of hosts and vectors. The strain that blocks the flea increases in frequency with the abundance of the host and the vector. Other parameter values (see **Appendix** for more details about the simulation procedure): *N_H_* = 2(1 + *f*[*t*]), *N_F_* = 5(1 + *f*[*t*]), γ = 0.2, *θ* = 1, *σ* = 0.25, *δ* = 0.9, *m_H_* = 0.002, *m_F_* = 0.04, *α_H_* = 0.1, *α_U_* = 0.01, *α_B_* = 0.2, *β_H_* = 0.1, *β_P_* = 0.06, *β_U_* = 0.05, *s*_0_ = 0.5, 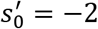.

## Discussion

The emergence and the evolution of plague results from a series of adaptations that increased the efficacy of flea-borne transmission of *Y. pestis* (3,4,7). But whether or not the blockage of the flea is an adaptation remains a controversial issue (7,14,15,17). Our analysis is an attempt to clarify the conditions that can promote the evolution of blockage. Here we consider a situation where a mutant bacteria with a distinct blockage strategy is introduced in a population of *Y. pestis* and we determine if such a mutant can invade or not. For instance, different genetic variants in the *hmsHFRS* operon are known to affect dramatically the colonization of the proventriculus and the formation of a biofilm: the *hmsFRS+* mutant is known to yield flea blockage while *hmsFRS*-never blocks the fleas and the mortality of fleas blocked by the *hmsFRS+* mutant is considerably larger than unblocked fleas (7,11). Does the gain in transmission due to blockage compensates this increased mortality? Our analysis allows us to answer this question. More specifically, the condition (5) shows that blockage is adaptive, in the absence of within-flea competition, if the ratio of mortality rates between blocked and unblocked fleas is lower than the ratio of transmission rates between blocked and unblocked fleas:

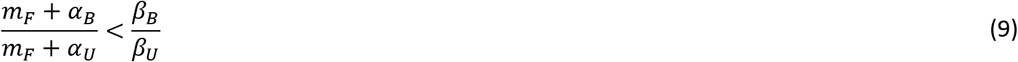

Available data on blocked and unblocked rat flea *Xenopsylla cheopsis* (one of the main flea vector) suggest that that the life expectancy of a blocked flea is around 2 days while the life expectancy of an infected (but unblocked) flea is around 100 days (11,19,32,7). The ratio between mortality rates of blocked and unblocked fleas is thus expected to be around 50. In other words, condition (9) indicates that transmission rate of blocked fleas must be 50 times higher than transmission rate or unblocked fleas for blockage to be adaptive. Available experimental data on *X. cheopsis* suggests that transmission of blocked fleas is likely to be much higher than this threshold value. First, the ratio of the biting rates of blocked and unblocked fleas is likely to be higher than 3 (19). Second, the number of *Y. pestis* bacteria transmitted by blocked fleas is several order of magnitudes higher (19). Given that regurgitation of a larger inoculum increases the chance of the bacteria to establish a successful infection in the mammalian host, the ratio 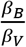 is likely to be higher than a few hundreds. Obviously, obtaining more accurate estimates of transmission and mortality rates in *X. cheopsis* (but also in other flea species) is particularly important to conclude on the adaptive nature of blockage.

Our analysis introduces also the possibility of within-flea competition between different variants of *Y. pestis*. In particular, we contend that the production of a biofilm may be a way to outcompete other bacteria in the foregut of the flea. Within-flea competition adds another dimension in the adaptive value of blockage. In particular, conditions (5) and (6) indicate that this mechanism is likely to promote the evolution of blockage. Recent experimental studies have explored the outcome of competition between different strains of *Y. pestis* in fleas (33–36). Unfortunately, experiments following the competition taking place between *hms* variants in the flea remain to be carried out.

Empirical evidence of plague dynamics reveal the highly epidemic nature of plague outbreaks which is likely to be driven by seasonal variations of the environment (26,27). In such a fluctuating environment, our analysis reveals that selection for blockage is likely to vary through time. Blockage should be more strongly selected at the onset of epidemics, when many hosts are uninfected. In contrast, blockage is expected to decrease when the epidemic is fading away because a smaller number of susceptible hosts are available. It would be interesting to study the variability of the ability to produce a biofilm and to block the fleas in natural populations. Analysis of bacteria sampled at different points in space or in time would allow to test our prediction that temporal fluctuations in the environment drives the maintenance of variability in *Y. pestis* populations.

Even though our model tries to capture multiple routes of transmission, it is important to acknowledge that plague transmission involves a multitude of host species (37). Our model focuses on a simple scenario with a single species of vertebrate host and a single species of flea. Yet, the competence of fleas, their propensity to develop blockage and their mortality rates (after blockage) are known to differ widely (7,32,38). Besides, the infectious blood source is also known to affect the development of *Y. pestis* in the fleas (39). A full understanding of the ecology and evolution of the plague thus requires a more comprehensive description of the network of host and vector species involved in its transmission.

## Acknowledgements

We thank Boris Schmid for very useful discussion and comments on an earlier version of this manuscript.

## Appendix

### Derivation of *R*_0_

The ability of the pathogen to invade an uninfected host population is determined by, *R*_0_, its basic reproduction ratio. To derive *R*_0_ we need to consider the dynamics of equation (1) when *S* = *N_H_* and *F_S_* = *N_F_*:

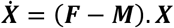

where:

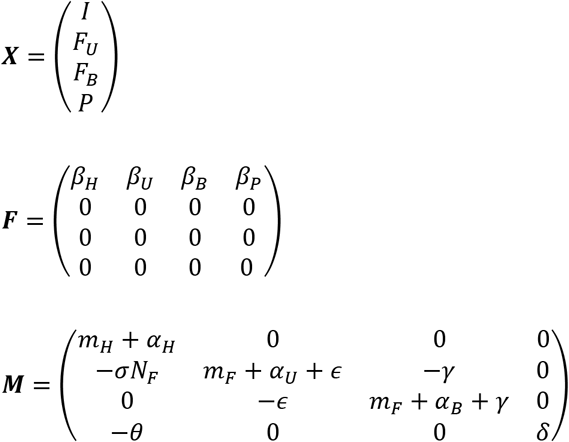

The basic reproduction ratio is the dominant eigenvalue of ***F.M*^−1^** which yields equation (3) in the main text.

### Pathogen evolution

To study pathogen evolution we first track the dynamics of a rare mutant invading the population of a resident pathogen when the system has reached and endemic equilibrium. For the sake of simplicity, we assume that coinfections with the resident and the mutant pathogens are not feasible but we do allow for superinfections in the vector which yields the dynamical system (1). In matrix form this yields the following dynamical system:

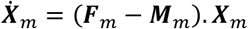

where:

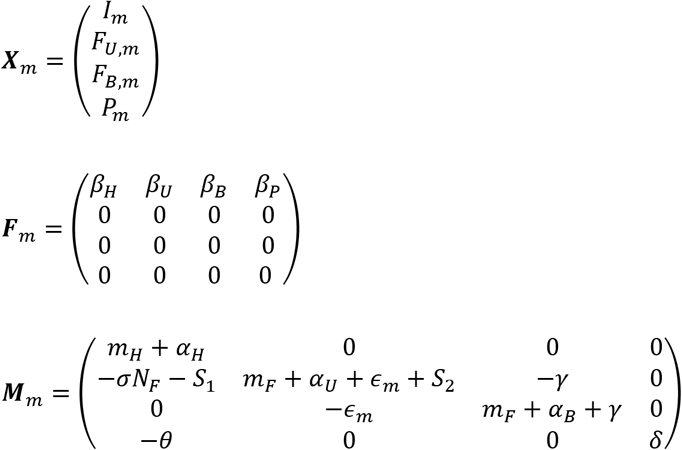

With *S*_1_ = *s*[*ϵ, ϵ_m_*]*Ī* and 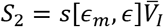

The basic reproduction ratio is the dominant eigenvalue of ***F**_m_.**M**_m_*^−**1**^ which yields equation (4) in the main text.

### Simulations

In **Figure 2** we present a simulation of the dynamical system (1) with two strains: one strain never blocks the flea (*ϵ*_1_ = 0) and another strain can block infected fleas (*ϵ*_2_ = 1). To illustrate the dynamics occurring during an epidemic we assumed that none of the vectors are initially infected (*F_S_* = *N_F_*) and we introduced a small density of infected hosts: *I*_1_ = 10^−4^ and *I*_2_ = 10^−6^. **Figure 2** shows the epidemiological and the evolutionary dynamics when condition (5) is satisfied or not (panel (B) and (A), respectively).

In **Figure 3** we present a simulation of the dynamical system (1) under the assumption that *N_H_* and *N_F_* vary periodically because of seasonality with two strains: one strain never blocks the flea (*ϵ*_1_ = 0) and another strain can block infected fleas (*ϵ*_2_ = 0.8). Under the parameter values we chose, the two strains can coexist in the long-term. We show the epidemiological and evolutionary dynamics for 3 consecutive seasons, when the system has reached a stable limit cycle.

---

**Figure S1:**
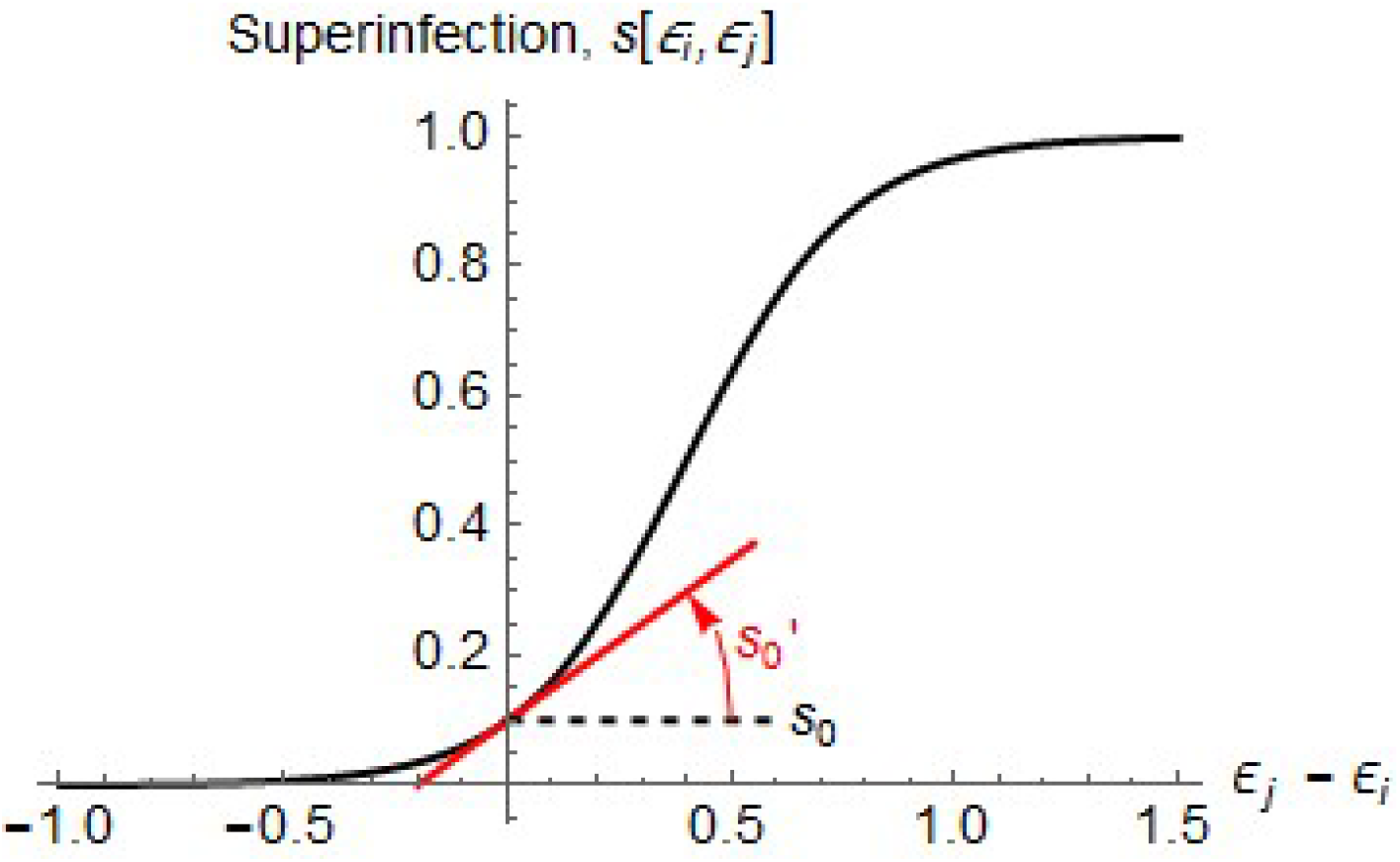
The superinfection function. This function measures the probability that a strain *j* with the blockage strategy *ϵ_j_*, outcompetes a resident strain *i* with the blockage strategy *ϵ_i_*, in the flea vector (see description of the parameters of the superinfection function in the main text). In the scenario illustrated in the above figure, higher investment in the production of a biofilm is associated with a within-host fitness advantage in the flea vector (*s*′_0_ >0).

**Figure S2:**
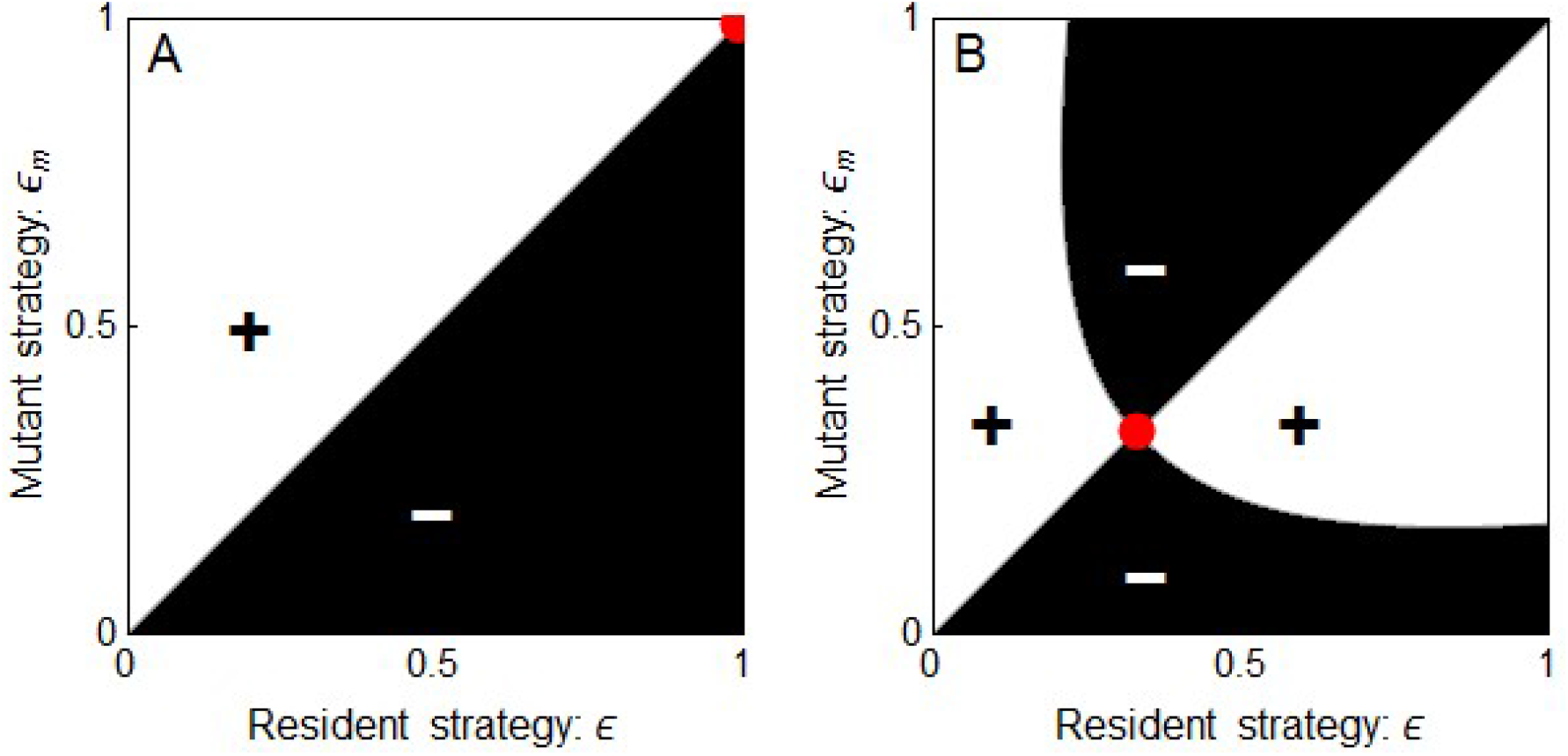
Pairwise invasibility plot on blockage. We use equation (4) to plot the ability of the mutant strategy *ϵ_m_* to invade a resident population with strategy *ϵ*. When *R_m_* > 1 the mutant can invade (white) and when *R_m_* < 1 the mutant fails to invade the resident population (black). In (A) 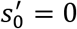 and in (B) 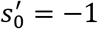. Pairwise invisibility plots can be used to find the ultimate evolutionary outcome (red dot) but also to identify pairs of strategies that can coexist. Panel (B) shows that an intermediate strategy can be evolutionary stable. Other parameter values: *N_H_* = 1, *N_F_* = 6, *γ* = 0.23, *θ* = 5, *σ* = 0.28, *δ* = 0.75, *m_H_* = 0.002, *m_F_* = 0.04, *α_H_* = 0.8, *α_U_* = 0.01, *a_B_* = 0.2, *β_H_* = 0.1, *β_P_* = 0.073, *β_U_* = 0.02, *β_B_* = 0.2, *s*_0_ = 0.5.

**Table S1:** Definitions of the main parameters of the model

| Main parameters | Definitions |
| --- | --- |
| $N_H$ | Density of the vertebrate host |
| $N_F$ | Density of the flea vector |
| $S, I$ | Densities of uninfected and infected hosts |
| $F_S, F_U, F_B$ | Densities of uninfected, infected-unblocked and infected-blocked fleas |
| $P$ | Density of free-living *Y. pestis* bacteria in the soil |
| $\beta_P$ | Infectivity of free-living propagules for the host |
| $\beta_H$ | Direct transmission rate among hosts |
| $\beta_U$ | Infectivity of unblocked fleas |
| $\beta_B$ | Infectivity of blocked fleas |
| $m_H$ | Natural mortality rate of the host |
| $m_F$ | Natural mortality rate of the flea |
| $\delta$ | Mortality rate of free-living propagules |
| $\theta$ | Rate of production of free-living propagules by infected hosts |
| $\alpha_H$ | Virulence (mortality induced by the pathogen) in the host |
| $\alpha_U$ | Virulence (mortality induced by the pathogen) in the unblocked fleas |
| $\alpha_B$ | Virulence (mortality induced by the pathogen) in the blocked fleas |

